# Whisker deprivation triggers a distinct form of cortical homeostatic plasticity that is impaired in the *Fmr1* KO

**DOI:** 10.1101/2024.09.23.614487

**Authors:** Alishah Lakhani, Washington Huang, Chris C. Rodgers, Peter Wenner

## Abstract

Mouse models of Fragile X Syndrome (FXS) have demonstrated impairments in excitatory and inhibitory sensory-evoked neuronal firing. Homeostatic plasticity, which encompasses a set of mechanisms to stabilize baseline activity levels, does not compensate for these changes in activity. Previous work has shown that impairments in homeostatic plasticity are observed in FXS, including deficits in synaptic scaling and intrinsic excitability. Here, we aimed to examine how homeostatic plasticity is altered *in vivo* in an *Fmr1* KO mouse model following unilateral whisker deprivation (WD). We show that WD in the wild type leads to an increase in the proportion of L5/6 somatosensory neurons that are recruited, but this does not occur in the KO. In addition, we observed a change in the threshold of excitatory neurons at a later developmental stage in the KO. Compromised homeostatic plasticity in development could influence sensory processing and long-term cortical organization.

## INTRODUCTION

Fragile X Syndrome (FXS) is a neurodevelopmental disorder that is the most common monogenetic form of intellectual disability, with up to 50% of FXS male patients receiving a co-diagnosis of autism spectrum disorder (ASD)^1^. In addition to cognitive impairments, patients with FXS can have seizures, circadian rhythm disruptions, and sensory hypersensitivity ^2,3^. FXS is caused by a trinucleotide repeat expansion that inactivates the *Fmr1* gene on the X-chromosome, resulting in the absence of fragile X messenger ribonucleoprotein (FMRP)^4^. FMRP is an mRNA translational repressor that plays an important role in synaptic function and plasticity ^3,5^. For instance, the mGluR (metabotropic glutamate receptors) theory of FXS states that the loss of FMRP increases long-term depression (LTD), a protein-synthesis dependent form of plasticity ^6,7^.

The *Fmr1* KO mouse cortex exhibits weaker whisker-evoked activity compared to the wild type (WT) ^8^. For instance, layer 2/3 (L2/3) excitatory and inhibitory neurons in the somatosensory cortex (S1) fire less frequently following whisker stimulation compared to WT neurons. Homeostatic plasticity encompasses a set of mechanisms thought to maintain activity levels within appropriate boundaries ^9,10^. When perturbations to cellular or network activity occur, compensatory changes in synaptic strength (homeostatic synaptic plasticity) and/or intrinsic membrane excitability (homeostatic intrinsic plasticity) are engaged to stabilize neural function. It is unclear why homeostatic plasticity does not compensate for the reduced whisker-evoked cortical activity in FXS mice. However, previous studies have suggested that both homeostatic synaptic plasticity ^11–14^, and homeostatic intrinsic plasticity can be impaired in *Fmr1* KO mice ^15^. Since this work has been largely performed in cultures or slices, we aimed to test the homeostatic capacity of FXS mice in an intact, *in vivo* system.

We chose to examine the barrel cortex for several reasons. The somatosensory cortex (S1) somatotopically represents the whiskers on the snout of a rodent; each individual whisker is represented in the contralateral cortex by a distinct region called the barrel ^16,17^. The whisker system is crucial for a mouse’s sensory experience, helping them navigate and interact with the environment. Individual whiskers can be stimulated to examine the responsiveness of cortical neurons in the corresponding barrel to a relevant sensory input, as the previous study showing weakened responses in the *Fmr1* KO had exploited ^8^. Unilateral whisker deprivation, removing all whiskers on one side of the snout, has been shown to trigger a homeostatic increase in the responsiveness of S1 cortical neurons to whisker stimulation ^18^. Finally, impairments in whisker-evoked processing are prevalent in different models of ASD in developing rodents ^8,19–23^. Thus, in order to investigate the homeostatic capacity of the *Fmr1* somatosensory cortex, we performed unilateral whisker deprivation during a critical developmental window (P14-P21, at the onset of whisking) to test homeostatic plasticity in the output layer of S1 (L5/6). We found reduced recruitment of L5/6 neurons during whisker stimulation in the KO at P16, but not P21. Further, we found that recruitment of this L5/6 cortical population was significantly increased following both 2- and 7-day whisker deprivation in the WT cortex, but not in the KO cortex. Our results show that while the KO L5/6 mouse cortex has some homeostatic capacity, it fails to compensate for reduced sensory input by recruiting a larger portion of the L5/6 population.

## RESULTS

### P16 WT RS neurons respond to whisker stimulation more than KO RS neurons

In order to compare excitability in the WT and *Fmr1* KO S1 cortex, we examined the regular spiking (RS, presumed excitatory) L5/6 neurons, as these layers provide the output of the cortical column, projecting to the secondary somatosensory cortex, motor cortex, subcortical structures, and the corticothalamic pathway ^17,24^. Since FXS and autism are neurodevelopmental disorders, we examined evoked responses during an early developmental stage where homeostatic plasticity mechanisms are most robustly expressed ^10,23,25^. Mice were lightly anesthetized at postnatal day 16 (P16), and a 64-channel probe was inserted into the barrel cortex to record spiking activity. We stimulated 9 individual whiskers at varying velocities to generate a velocity response curve (VRC) that measured the responsiveness of cortical L5/6 RS neurons, and curated units with Kilosort 2.5 (see methods).

We first tested whether there were any differences in spontaneous activity in WT and KO L5/6 excitatory neurons at baseline (no whisker stimulation), and observed that the spontaneous firing rate of individual neurons was no different (Figure 2A). We then calculated the whisker-evoked VRC of the columnar whisker (CW, also referred to as the principal whisker), histologically identified following the experiment (Figure 1B, see methods). The average CW VRC of KO L5/6 neurons was reduced compared to the WT, demonstrating that these neurons were less responsive to whisker stimulation (Figure 2B). These VRCs will be referred to as overall VRCs as they include all L5/6 RS neurons, whether they were responsive to the CW stimulation or not. Therefore, this population response could be due to decreased sensitivity at the single neuron level, decreased proportion of responsive neurons, or both. We found that there was no difference between WT and KO VRCs if we only included neurons that significantly respond to CW stimulation (CW-responsive neurons; Figure 2C), suggesting that there was a reduction in the proportion of CW-responsive L5/6 neurons. Indeed, we found a reduced proportion of neurons that were CW-responsive in the KO, albeit this did not reach significance (Figure 2D). In fact, when we compared the proportions of all whisker-responsive L5/6 neurons in WT and KO mice (proportion of neurons responsive to any of the 9 whiskers stimulated compared to total number of neurons – whisker responsive or not), we found that there was a significant decrease in the KO (Supplemental Figure 1A).

**Figure 1.**
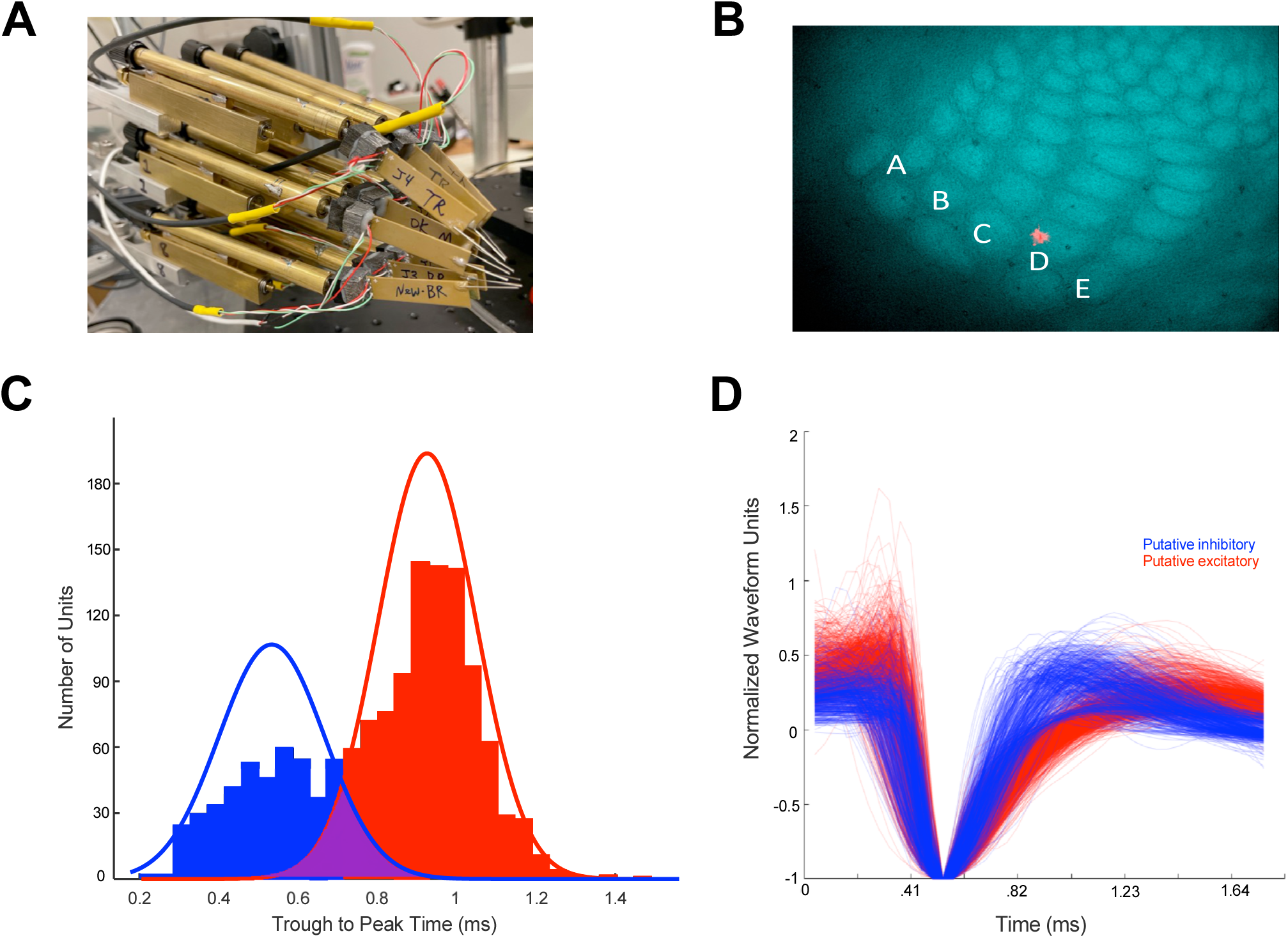
Classification of single units. A) Nine whiskers were inserted into a 3×3 piezoelectric stimulator array, ideally inserting the whisker that elicited the strongest response across all 64 channels in the middle of the array. B) Example histology section of the barrel cortex. Red fluorescence indicates the location of the probe (in this case, D1). C) The bimodal distribution of waveforms from extracellular recordings using the trough to peak time. A Gaussian Mixture Model was fit to this distribution. The point at which the two curves intersect was used as the boundary between putative inhibitory and excitatory units. Overlap area is approximately 10%. D) Mean waveform of all units used in the analyses. Blue waveforms represent putative inhibitory neurons, and red waveforms represent putative excitatory neurons.

**Figure 2.**
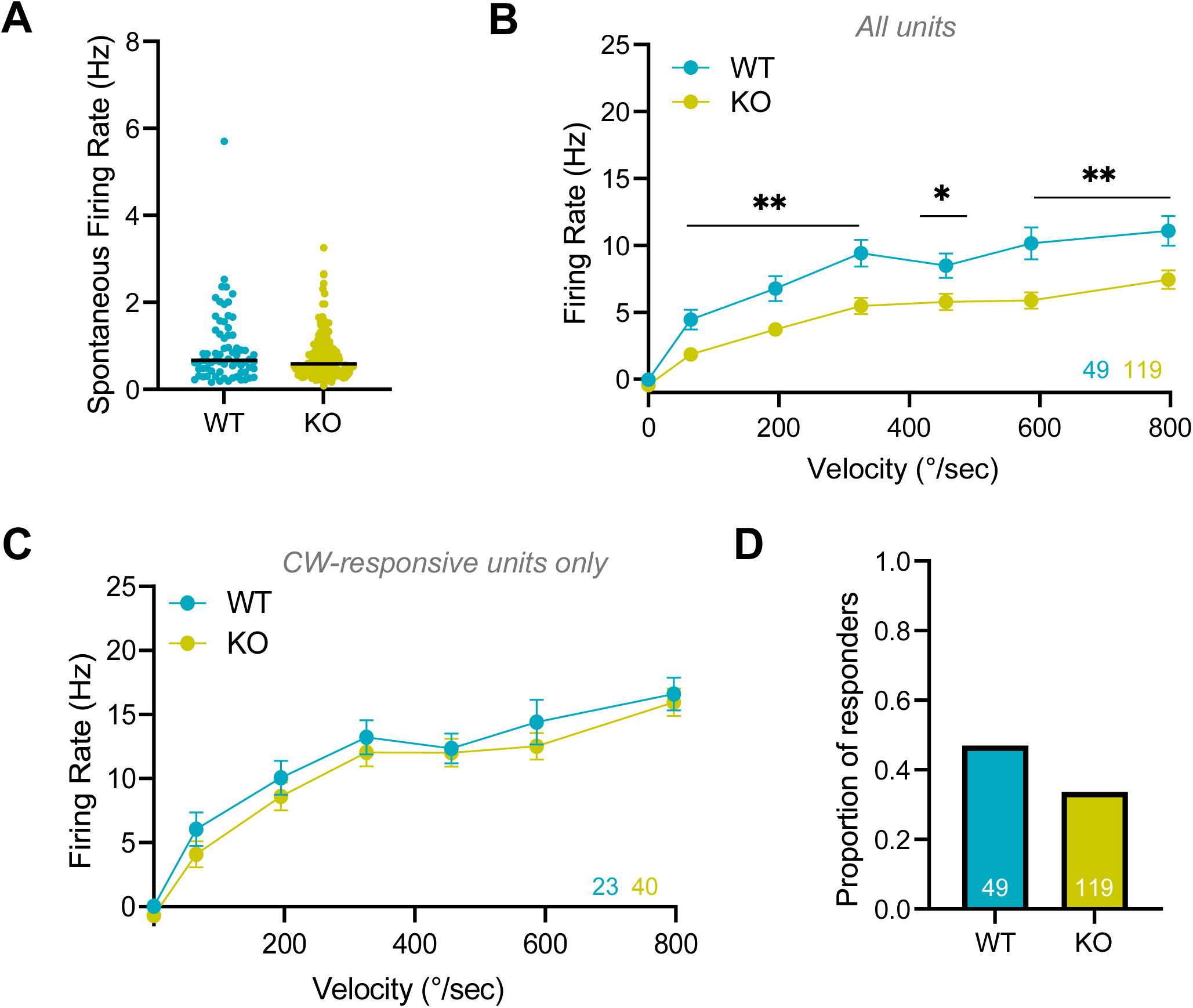
Baseline WT and *Fmr1* KO whisker-responsiveness at P16. A) Spontaneous firing rates of WT and *Fmr1* KO neurons. B) Overall velocity response curve (VRC) of all neurons following columnar whisker (CW) stimulation at varying velocities. Number of units for each condition is color-coded and shown at the bottom-right. C) VRC of CW-responsive neurons only. D) The proportion of neurons that significantly respond to CW stimulation. * p < 0.05, ** p < 0.01.

We were also interested in evaluating the VRC for the best whisker (BW) of the neuron, which is the whisker that elicited the strongest response for that neuron, regardless of the anatomical location of the cell (including neurons in barrels and septa). Like the VRC for CW-responsive neurons, the average BW VRC was nearly identical in WT and *Fmr1* KO L5/6 RS neurons (Supplemental Figure 2A). These results suggested that at baseline, there were no differences in the excitability of whisker-responsive (CW or BW) neurons in WT and KO neurons. Thus, while the proportion of neurons recruited in a column was reduced in the KO, the whisker-responsive neurons were similar. Since the VRCs produced by stimulating the CW or BW were similar, for clarity we will discuss CW VRCs in all further comparisons, but show BW VRCs in Supplemental Figure 2.

### P16 2-day whisker-deprived WT and KO neurons increased their responsiveness, but only WT neurons increased neuronal recruitment

In order to test the homeostatic capacity of RS neurons, we unilaterally trimmed whiskers to 2mm at P14, and recorded activity from the barrel cortex at P16 (2 days of whisker deprivation – WD). We found that following WD, there was an increase in overall VRCs from CW stimulation across all velocities (Figure 3B). We also found a slight increase in the VRC of CW-responsive neurons (Figure 3C). In this case, the main increase in the VRC was due to an increased proportion of neurons within a barrel that significantly responded to the CW deflection (Figure 3D, ∼45% to 80% after WD). Therefore, following sensory deprivation, more of the WT S1 L5/6 RS neurons were recruited in a homeostatic manner, and of those that responded, there was a slight increase in the evoked output.

**Figure 3.**
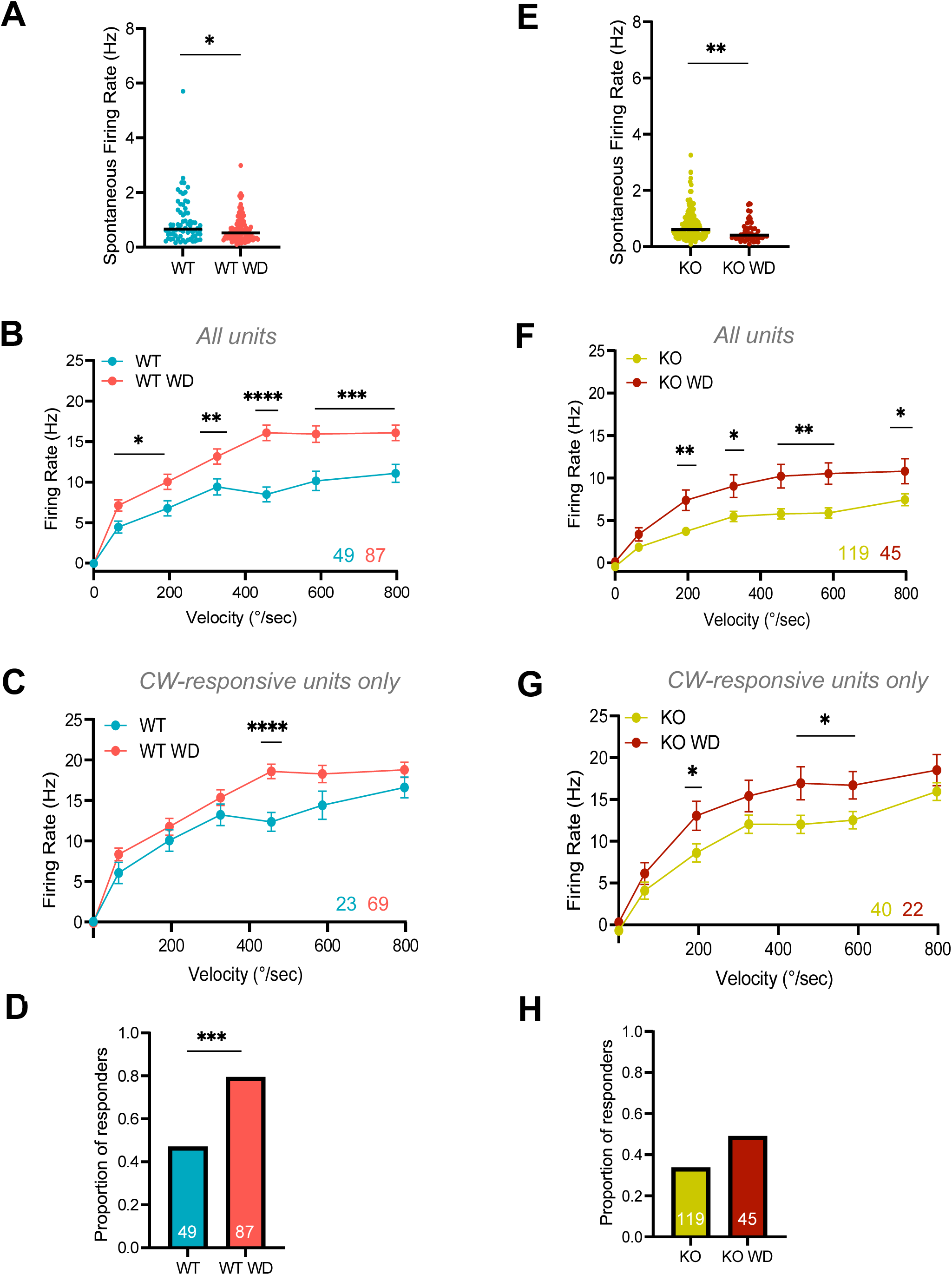
WT and *Fmr1* KO whisker-responsiveness following whisker deprivation at P16. A) Spontaneous firing rates of WT and WT whisker-deprived (WT WD) neurons. B) Overall velocity response curve (VRC) of all WT and WT WD neurons following columnar whisker (CW) stimulation at varying velocities. Number of units for each condition is color-coded and shown at the bottom-right. C) VRC of CW-responsive neurons only for WT and WT WD. D) The proportion of neurons that significantly respond to CW stimulation. E) Spontaneous firing rates of KO and KO whisker-deprived (KO WD) neurons. F) Overall velocity response curve (VRC) of all KO and KO WD neurons following CW stimulation at varying velocities. G) VRC of CW-responsive neurons only for KO and KO WD. H) The proportion of neurons that significantly respond to CW stimulation. * p < 0.05, ** p < 0.01, *** p < 0.001, **** p < 0.0001.

We had hypothesized that KO neurons would demonstrate some impairment in homeostatic plasticity. CW stimulation shifted the overall VRC significantly upward following 2-day sensory deprivation in the KO, for all neurons and for CW-responsive neurons alone (Figure 3F and 3G). However, the proportion of neurons that responded to CW stimulation in the KO did not increase significantly with WD (34% to 49% after WD, Figure 3H). The findings suggest that while *Fmr1* KO L5/6 neurons at P16 demonstrate some homeostatic capacity following WD, there is an impairment in the compensatory increase of the population recruited by CW stimulation, which is considerably reduced compared to the WT WD neurons (49% vs 80%). Interestingly, the spontaneous firing rate following WD in both the WT and KO conditions decreased compared to their respective baseline values (Figures 3A and 3E).

### P21 KO RS neurons exhibit an increased threshold to whisker stimulation compared to WT neurons

We extended these studies to P21 with a 7-day whisker deprivation, but will first compare the P21 WT and KO at baseline (without whisker deprivation). We first tested whether there were any changes in spontaneous activity between the WT and KO L5/6 RS neurons. We found that, like P16, the spontaneous firing rate of single neurons was the same in both genotypes (Figure 4A). We then examined the overall VRCs from CW stimulation, and interestingly, discovered that the KO VRC exhibited a shift in the shape of the curve compared to the WT VRC (Figure 4B). This trend was even more apparent when we analyzed the VRC using CW-responsive neurons, where the lowest velocity whisker deflection to elicit a response was higher in the KO than in the WT (Figure 4C). These results suggested a potential change in the sensitivity or threshold of the KO L5/6 RS neurons. Indeed, when we calculated the minimum whisker stimulation velocity that neurons required to elicit a significant whisker-evoked response, we found that this velocity was greater in the KO condition (Table 1). However, at the faster whisker stimulations, we observed no difference in the firing rate of KO neurons, indicating that the decrease in spiking activity only occurred at slower velocities. Moreover, there was no significant difference in the proportion of neurons that responded to CW stimulation at the fastest velocity (Figure 4D). These results demonstrate an increase in the threshold to activate KO L5/6 RS neurons at P21, but no change in the number of cells recruited. Interestingly, this decreased sensitivity is unique to P21 neurons, and this change occurred in the responsiveness of KO L5/6 RS neurons in just five days.

**Figure 4.**
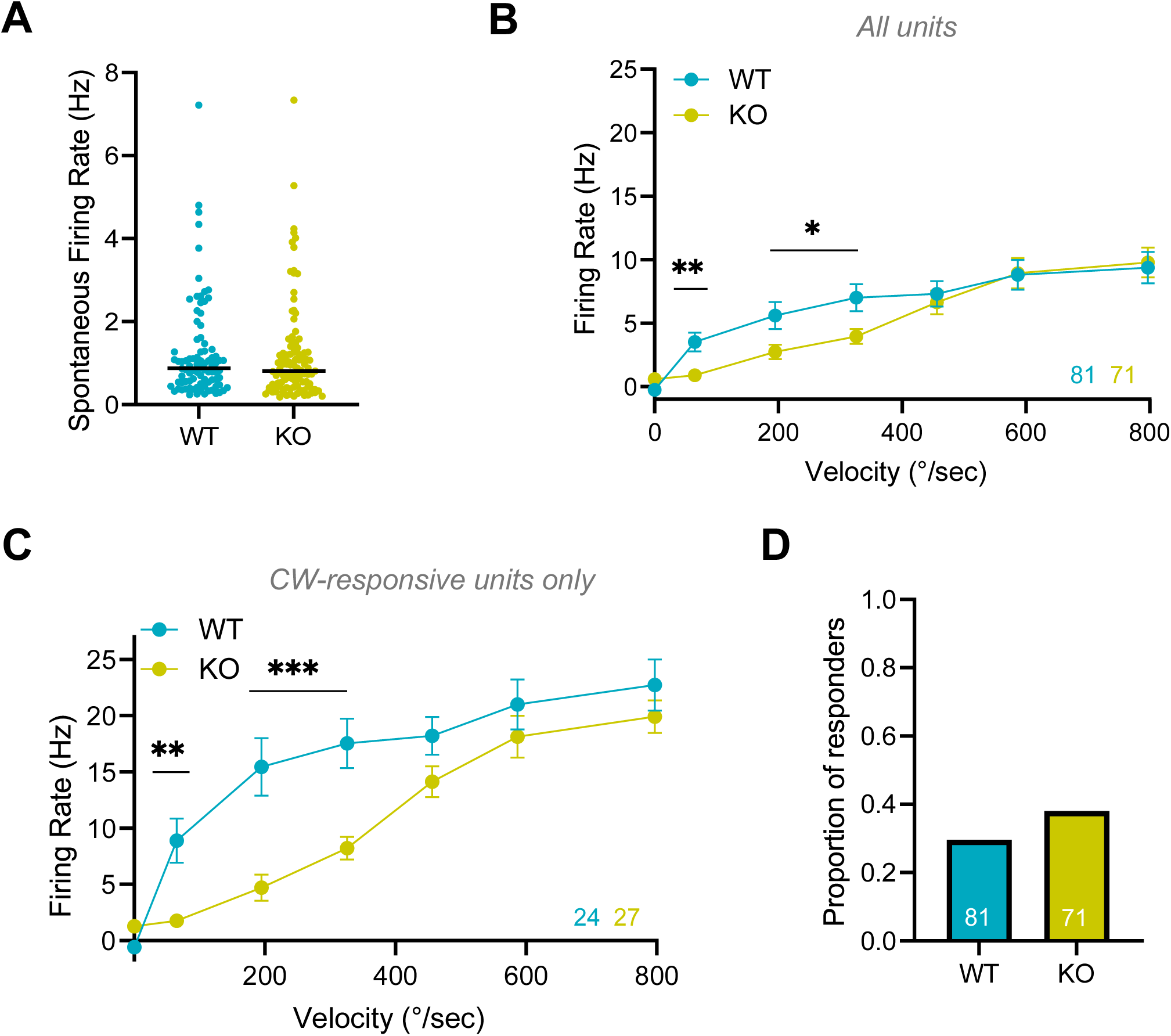
Baseline WT and *Fmr1* KO whisker-responsiveness at P21. A) Spontaneous firing rates of WT and *Fmr1* KO neurons. B) Overall velocity response curve (VRC) of all neurons following columnar whisker (CW) stimulation at varying velocities. Number of units for each condition is color-coded and shown at the bottom-right. C) VRC of CW-responsive neurons only. D) The proportion of neurons that significantly respond to CW stimulation. * p < 0.05, ** p < 0.01, *** p < 0.001.

**Table 1.**
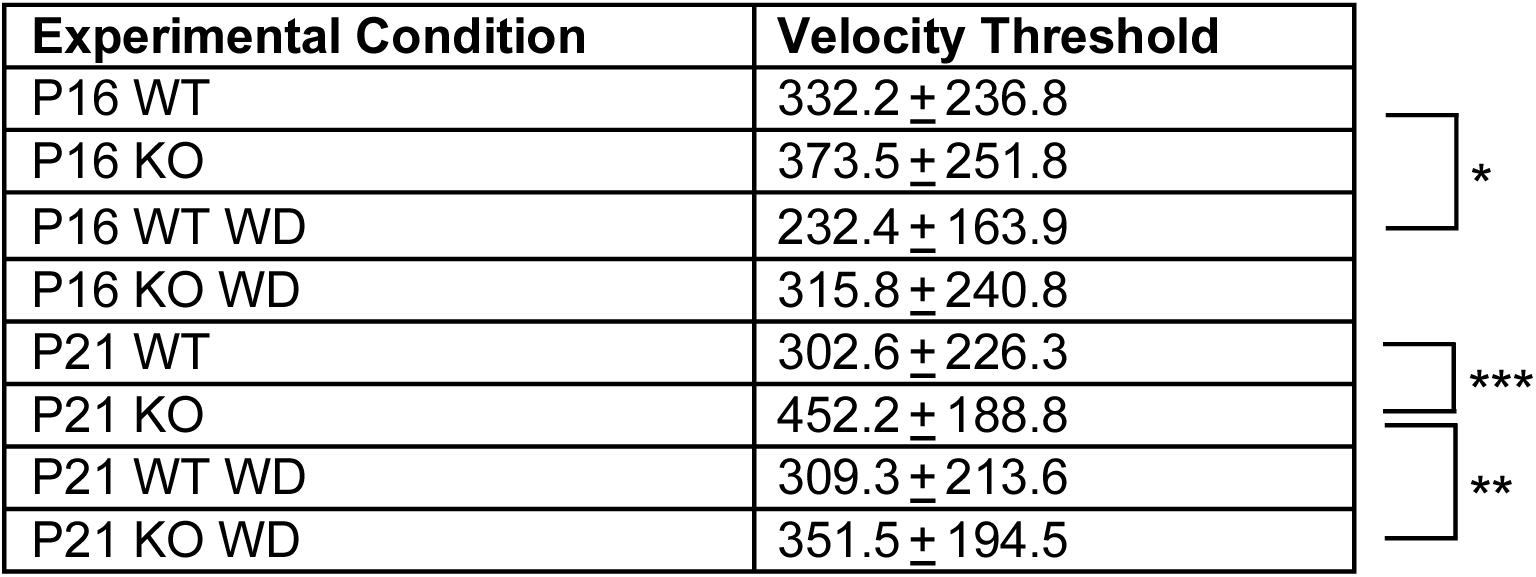
Velocity threshold for each condition, calculated using only CW-responsive neurons. All values are mean ± standard deviation. * p < 0.05, ** p < 0.01, *** p < 0.001.

### P21 7-day whisker-deprived WT and KO neurons increased their responsiveness, but only WT neurons increased neuronal recruitment

Next, we examined the effect of a longer (7-day) sensory deprivation period on WT L5/6 RS neurons. Whiskers were unilaterally trimmed to 2mm at P14 and then trimmed every other day until the day of recording, P21. We found that following whisker deprivation, there was a slight increase in the overall VRC in response to CW stimulation (Figure 5B). VRCs of CW-responsive neurons were no different in WT and whisker-deprived WT mice (Figure 5C). Instead, the increase in the overall CW VRC was a direct result of a strong increase in the proportion of responsive neurons (30% to 57% after WD, Figure 5D). Thus, a 7-day whisker deprivation in WT mice did not alter the responsiveness of L5/6 RS neurons that responded, but the network was able to compensate by recruiting twice as many neurons.

**Figure 5.**
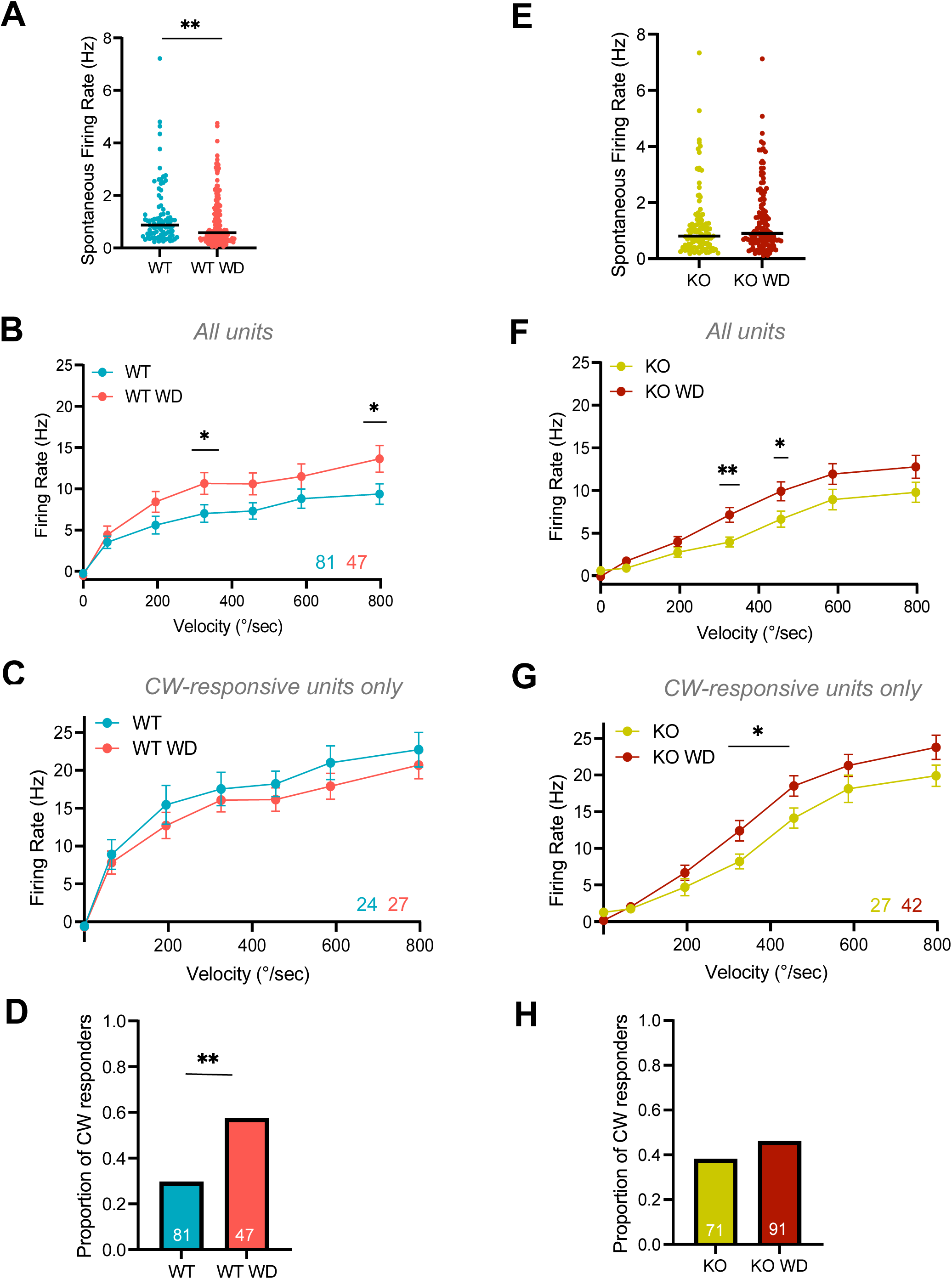
WT and *Fmr1* KO whisker-responsiveness following whisker deprivation at P21. A) Spontaneous firing rates of WT and WT whisker-deprived (WT WD) neurons. B) Overall velocity response curve (VRC) of all WT and WT WD neurons following columnar whisker (CW) stimulation at varying velocities. Number of units for each condition is color-coded and shown at the bottom-right. C) VRC of CW-responsive neurons only for WT and WT WD. D) The proportion of neurons that significantly respond to CW stimulation. E) Spontaneous firing rates of KO and KO whisker-deprived (KO WD) neurons. F) Overall velocity response curve (VRC) of all KO and KO WD neurons following CW stimulation at varying velocities. G) VRC of CW-responsive neurons only for KO and KO WD. H) The proportion of neurons that significantly respond to CW stimulation. * p < 0.05, ** p < 0.01.

Since the spiking activity of KO L5/6 RS neurons was already different than the WT neurons at baseline across the lower whisker stimulation velocities, we were curious to see how whisker deprivation would affect responsiveness and recruitment. The overall CW VRC of KO neurons following sensory deprivation was higher than without WD, demonstrating that these neurons homeostatically became more responsive to whisker stimulation (Figure 5F). We discovered the same result when examining only the VRCs of CW-responsive neurons (Figure 5G). Interestingly, as in P16 KO neurons, the proportion of CW-responsive neurons was not significantly different following deprivation in the KO at P21 (Figure 5H). Thus, the increased VRC was directly due to an increase in the responsiveness of individual neurons that were CW-responsive. Therefore, although the spiking activity increased in both WT and KO neurons after a sensory perturbation, the recruitment of neurons in the cortical column was different: only in the WT did the proportion of recruited cells significantly increase. Further, although WD increased the overall VRC in the KO, these mice were still less responsive than the WT at the lowest frequencies. Finally, following 7-day whisker deprivation, the spontaneous firing rate of WD neurons in the WT significantly decreased, but remained unchanged in the KO at P21 (Figure 5A and 5E).

### WT and KO FS neurons respond similarly to RS neurons at P16

Finally, we wanted to test whether there were changes in fast-spiking (FS) putative inhibitory neurons for each condition and how they might impact what we observed in the RS neurons. Unfortunately, we were not able to record from a sufficient number of FS units at P21 or in CW-responsive neurons at P16, but here, we present overall VRCs and proportions of responders for FS neurons at P16. At baseline, without whisker deprivation, we observed a weakening in the overall CW VRC of KO FS neurons compared to WT FS neurons (Figure 6A). This was, at least in part, due to a slight decrease in the proportion of FS neurons that were recruited following CW stimulation (Figure 6B). Following a 2-day whisker deprivation in WT mice, we observed an increase in the overall VRC, and this was due to an increase in the proportion of neurons recruited in WT mice (Figure 6C and 6D). In the KO mice, we observed an increase in the overall CW VRC after WD, but observed no change in the proportion of CW-responsive neurons (Figure 6E and 6F). These results are very similar to what we observed for L5/6 RS neurons at P16, suggesting that both excitatory and inhibitory neurons might respond similarly to whisker deprivation.

**Figure 6.**
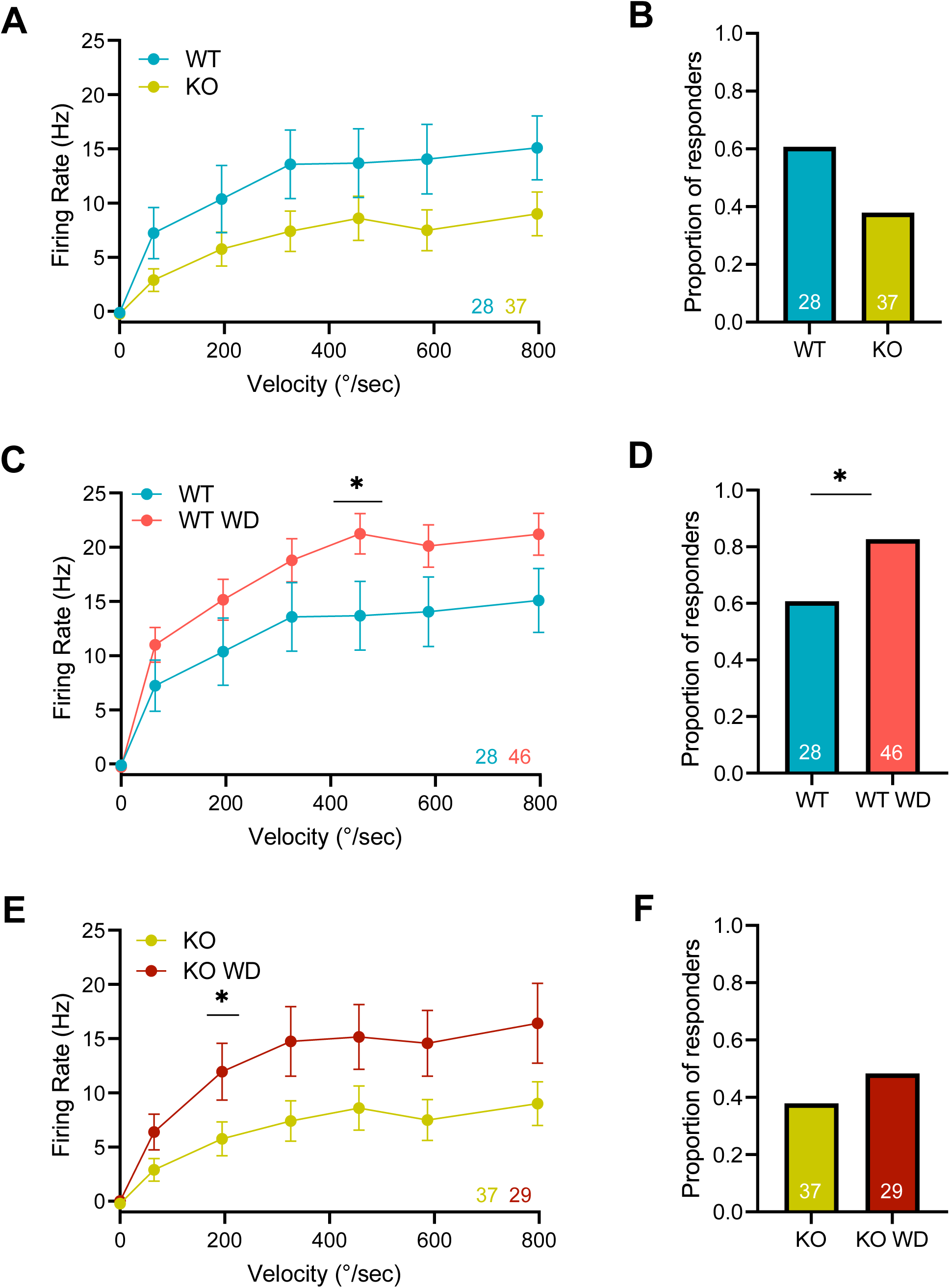
WT and *Fmr1* KO FS whisker responsiveness at baseline and following whisker deprivation at P16. A) Overall velocity response curve (VRC) of all neurons following columnar whisker (CW) stimulation at varying velocities for WT and KO mice at baseline. Numbers of units for each condition is color-coded and shown at the bottom-right. B) The proportion of neurons that significantly respond to CW stimulation. C and D) Similar to A and B except for WT and WT WD (whisker-deprived) neurons. E and F) Similar to A and B except for KO and KO WD (whisker-deprived) neurons. Number of units for each condition is color-coded and shown at the bottom-right. * p < 0.05.

## DISCUSSION

In this study, we have asked about the homeostatic capacity of WT and *Fmr1* KO L5/6 neurons, as this represents the output of the cortical column in the barrel cortex and therefore, the drive to downstream brain regions in the sensory circuit. We observe homeostatic increases in whisker-evoked responses in this part of the barrel cortex in WT mice following whisker deprivation. Further, we show that the KO can express homeostatic plasticity in some ways, but fails in other ways. The overall L5/6 whisker-evoked response exhibited a compensatory increase following WD in both the WT and KO mice. However, unlike WT littermates, following WD, the KO failed to recruit larger proportions of the L5/6 population with whisker stimulation. In addition, at baseline, overall whisker-evoked responses were weaker in the KO compared to WT, although this occurred through different mechanisms at the two developmental stages. These results suggest that the output of the S1 cortex is altered in FXS mice at baseline and in their homeostatic capacity.

### Baseline differences in WT and KO neurons

Changes in the overall VRCs can occur because of changes in the proportion of neurons that are recruited by the evoked synaptic activity (proportion of responsive neurons) and/or changes in the responsiveness of neurons that already respond to whisker stimulation. At P16, we observed that overall VRCs were reduced for L5/6 KO RS neurons (Figure 2B). A prior study had shown that KO L2/3 RS neurons demonstrated a decrease in the whisker-evoked overall VRC, in anesthetized adult and juvenile *Fmr1* KO mice, as well as awake adult mice ^8^. We further examined what may have caused such a reduction in the overall VRC in our study, and found that it was due to a reduction in the proportion of L5/6 excitatory neurons that were recruited following CW stimulation in the KO mice compared to the WT mice (Figure 2D). Similar to these results, previous work had demonstrated that fewer L2/3 RS neurons responded to whisker stimulation in juvenile *Fmr1* KO mice ^19^. On the other hand, the CW-responsive VRCs were no different in the WT and KO mice (Figure 2C). Therefore, although fewer L5/6 RS neurons were recruited by whisker stimulation in the KO, the ones that were recruited responded to the same extent as the WT. The similarity of the VRCs of the whisker-responsive units in the WT and KO could be due to a compensatory increase in intrinsic excitability; increased membrane excitability has been observed repeatedly in *Fmr1* KO mice in layers 2/3, 4, and 5 ^8,20,26,27^.

We observed that L5/6 FS neurons had a reduced overall VRC to CW stimulation, albeit not significant (Figure 6A). Similar to the L5/6 RS neuronal changes, this reduced VRC appears to be mediated by a reduction in the proportion of FS neurons that were recruited. Previous work has suggested that L2/3 parvalbumin (PV) neurons are less responsive to sensory stimulation in the KO ^8,28,29^. This result is plausible given the significant connectivity between RS and FS neurons; thus, if fewer excitatory neurons are recruited, fewer inhibitory neurons may also be recruited. Further investigation of changes in the GABAergic circuitry and therapeutic strategies to rescue inhibitory neuron deficits may prove to be useful in combating FXS circuit deficits ^30^.

Later in development, at P21, we also observed a reduction in the overall VRC of the KO compared to the WT, but not in the same way. First, the proportion of CW-responsive neurons was no different in the KO, so whisker stimulation was capable of recruiting the same number of L5/6 RS neurons (Figures 4D). This was consistent with a previous study that found no change in the proportion of whisker-responsive L2/3 neurons in the adult *Fmr1* KO mouse ^8^. However, while the CW-responsive L5/6 RS neurons demonstrated weaker whisker-evoked activity in the KO, this only existed in the lower half of the whisker stimulation velocities (Figure 4C). This would mean that the neurons would need a stronger signal from incoming connections to ensure a full neuronal response. Intriguingly, even though KO L5/6 RS neurons initially require a faster whisker stimulation to elicit a whisker-evoked response, the VRCs were similar in the WT and KO neurons at the three fastest whisker stimulation velocities. Thus, despite the altered sensitivity, once the KO neurons were sufficiently stimulated, they responded normally, possibly through homeostatic mechanisms. These alterations in response threshold could impact learning abilities, which are an important clinical phenotype in FXS patients.

Our results showing differences in the recruitment of neurons at P16 but not at P21 could imply that there are developmental changes that occur in the KO mouse model. There may be homeostatic compensatory mechanisms at play that allow for the recovery of some cortical function by P21. This result is consistent with previous findings demonstrating that early developmental deficits in cortical circuitry are eventually resolved, suggesting developmental delays in FXS mice ^3^. Examples of developmental delay in the *Fmr1* KO model include the synaptic strength of the L4 to L3 connection, the window for long-term potentiation, and the transition of GABA from depolarizing to hyperpolarizing currents ^21,23,31^.

### Plasticity in WT neurons

In WT mice, we observed that both 2- and 7-day unilateral whisker deprivation led to an increase in the overall VRC of L5/6 RS neurons (increase in whisker-evoked spiking). This is consistent with a previous study using unilateral whisker deprivation that showed increased spiking in L4 and L2/3 RS neurons following CW stimulation by 7-day WD ^18^. Several previous studies have found that removing one row of whiskers (typically the D-row) for 5-10 days leads to a weakening of the deprived CW-evoked spiking in excitatory neurons ^32–34^. However, D-row deprivation represents a competitive condition where the CW is deprived but adjacent whiskers can compete for cortical space and therefore can trigger Hebbian sorts of plasticity. On the other hand, unilateral whisker deprivation appears to trigger a homeostatic increase in RS spiking, as competing whiskers have been removed. Therefore, it is likely that selective whisker deprivation may favor Hebbian plasticity, while unilateral or complete whisker deprivation triggers a more homeostatic response. The increase in the overall VRC that we observed following both 2- and 7-day whisker deprivation in WT mice was due to a significant increase in the proportion of CW-responsive neurons (Figure 3D & 5D). How does this homeostatic increase in the recruitment of L5/6 RS neurons occur? Previous work has shown that unilateral whisker deprivation can produce alterations in the synaptic circuitry in L2/3 and L5 RS neurons ^35,36^. However, several other studies have demonstrated that such a deprivation leads to weakening of synaptic strength onto L2/3 and L5 RS neurons, which would appear to be anti-homeostatic ^18,32,37^. A likely mechanism contributing to the increased responsiveness to whisker stimulation is an increase in the excitability of the membrane (intrinsic excitability), which has been reported in both L2/3 and L5 RS neurons following unilateral WD ^38,39^. Increases in intrinsic excitability can explain the stronger sensory-evoked responses, but is surprising in terms of the observation that spontaneous activity of RS neurons is reduced after 2- and 7-day WD (Figure 3A & 5A). Reductions in spontaneous activity have also been described following complete visual deprivation in the monocular visual cortex ^40^.

Finally, we examined changes in L5/6 FS inhibitory neurons at P16 as well, and found increases in the overall VRCs following 2-day WD (Figure 6E). This occurred, at least in part, through increases in the proportion of recruited FS neurons (Figure 6F). Therefore, the increase in excitatory L5/6 neurons after 2-day WD was not due to reduced activity in inhibitory FS neurons. It will be important to identity the homeostatic mechanisms exhibited by FS neurons follow unilateral whisker deprivation. These findings suggest that RS and FS neurons may share some features of plasticity following WD at this age.

### Plasticity in KO neurons

The homeostatic capacity of *Fmr1* KO L5/6 neurons was markedly different than what was observed in WT neurons. Following 2-day WD, the overall VRC for L5/6 excitatory neurons did increase, but not to the same extent as WT neurons (Figure 3B vs 3F). This increase appears to occur largely through an increase in the CW-responsive VRC (Figure 3G). The inability of the KO WD VRC to achieve levels similar to the WT WD VRC is due to the fact that the proportion of recruited neurons does not increase following WD in the KO (Figure 3D & 3H). Therefore, 2-day WD in the KO leads to a compensatory increase in the response of CW-responsive neurons, but cannot recruit larger proportions of these neurons and therefore, the overall increase in the output of the cortical column is muted. Similar to our results for the L5/6 RS neurons, L5/6 FS neurons show the same deficits in the KO after WD (increase in the overall VRC but not in the proportion of CW-responsive neurons – Figure 6E & 6F).

Following 7-day WD (P21) in the KO, the overall VRC for L5/6 excitatory neurons did increase, but again, was not as effective as in the WT (Figure 5B vs 5F). While 7-day WD in the WT led to a significant increase in the proportion of CW-responsive neurons, again, this did not happen in the KO (Figure 5H). The increase that was observed in the overall VRC following 7-day WD in the KO appears to occur largely through an increase in the CW-responsive VRC (Figure 5G). Thus, as occurred following 2-day WD in the KO, the compensation continued to manifest predominantly as an increase in responsiveness among the neurons that were already responsive to whisker stimulation. If one of the mechanisms for increased responsiveness following unilateral WD is an increase in the intrinsic excitability, then our results could suggest that some neurons can increase excitability (CW-responsive), while others do not. This possibility is very similar to the impairments in homeostatic intrinsic plasticity that we previously observed in KO cortical cultures, where some cells increased their ability to fire action potentials while other cells did not ^15^. Another clear difference between 7-day whisker-deprived WT and KO L5/6 neurons was the shape of the VRC. As observed at baseline (without WD), the KO VRC after WD was clearly weaker at lower, but not higher velocities (Figure 5). Therefore, WD did not resolve this feature.

Our results show a deficit in the ability to increase neuronal recruitment in a compensatory fashion. In addition to perturbations in homeostatic intrinsic plasticity, it seems likely that impairments in synaptic plasticity contribute. *Fmr1* KO cortical neurons show impairments in activity-dependent spine dynamics ^3,41^. In fact, one particularly relevant study showed that WT L5 S1 cortical neurons alter the rate of synapse elimination following unilateral WD, but this plasticity is absent in the *Fmr1* KO ^42^. Regardless of the mechanism, reduced recruitment of KO L5/6 neurons could result in inappropriate sensory processing and output, as L5/6 neurons are critical for sending output to other cortical and subcortical regions. For instance, reduced signaling could adversely affect the precision and accuracy of processing tactile information from whiskers, potentially impairing the mouse’s ability to detect, discriminate, and respond to sensory stimuli ^14,43^. These sensory impairments are observed in both FXS animal models and patients.

Few studies have investigated homeostatic plasticity in the FXS model, but deficits have been reported. Previous work has demonstrated an exaggerated homeostatic intrinsic plasticity ^44^ and a failure of homeostatic synaptic scaling ^11,14^. Our lab previously demonstrated that homeostatic intrinsic plasticity is impaired in *Fmr1* KO cortical cultures ^15^. Our current findings of impaired homeostatic recruitment in the *Fmr1* KO cortex following a physiologically realistic sensory deprivation suggest homeostatic plasticity deficits in the intact system, and could explain some of the phenotypes associated with this neurodevelopmental disorder. A deeper understanding of these changes could offer insights into how early developmental abnormalities may influence long-term cortical organization and sensory processing impairments in neurodevelopmental disorders such as FXS and ASDs.

## ACKNOWLEDGEMENTS

We would like to thank Gary Bassell for helping start our mouse colony. We would also like to thank Bill Goolsby for his help with design and construction of equipment used in this study. Finally, we would like to thank Caleb Wright, Garrett Stanley, and Nicholas Au Yong for their advice on spike sorting and experimental methods. This research was supported by the following funding source from NINDS: R01NS065992.

## AUTHOR CONTRIBUTIONS

Conceptualization was by A.L and P.W. All experiments were carried out by A.L., and W.H. contributed to histology. A.L., C.C.R., and P.W. carried out data analysis and contributed to the writing of the manuscript.

## METHODS

### Mice

Heterozygous female *Fmr1* mice (X-linked gene; The Jackson Laboratory, Strain #003025, backcrossed on C57BL/6J background) were crossed with wild type (WT) C57BL/6J males (Jackson Laboratory) to generate litters of pups with mixed genotypes (*Fmr1* KO, *Fmr1* heterozygous, and WT mice). Genotyping was outsourced using Transnetyx, an automated genotyping PCR service, after validation with in-house PCR. For all experiments, *Fmr1* KO male pups were compared to WT male littermate controls. The mice were housed in a 12-hour light/dark cycle and the animal protocol was approved by the Institutional Animal Care and Use Committee at Emory University.

### Whisker deprivation

Mice were lightly anesthetized with isoflurane, and all the whiskers on the right side of the vibrissal pad were trimmed to approximately 2 mm at postnatal day 14 (P14). For experiments conducted at the 2-day time point (∼P16), this initial deprivation was the only occurrence. For experiments conducted following a 7-day WD (∼P21), whiskers on the right were trimmed every 48 hours, allowing for 48-72 hours to pass after the last trimming session in order to ensure whiskers have regrown sufficiently for whisker stimulation at P21 or P22. Mice that did not undergo whisker deprivation were still anesthetized with isoflurane on corresponding days, and had all the whiskers on the right side of the vibrissal pad acutely trimmed immediately before whisker stimulation on the day of the experiment. Left whiskers were never trimmed.

### Electrophysiology Recordings

Male mice (P16-P22) were anesthetized with isoflurane and chlorprothixene (0.02 mg dissolved in 10 mL saline, 1 mL injected intraperitoneally). Mice were transferred to a heating pad (37 °C), and the snout was inserted into a nose cone for consistent oxygen and isoflurane. Depth of anesthesia was monitored by breathing rate. A head plate was dental cemented onto the skull, and a small craniotomy was made over the barrel cortex – 3 mm lateral and 1.5 mm caudal to bregma. Anesthesia was then reduced (0.6-1% isoflurane) to maintain a respiratory rate of 1 breath/sec or slightly faster to obtain electrophysiology recordings.

Extracellular recordings were made using silicon probes (H3, Cambridge NeuroTech, United Kingdom) with 64 recording sites covering 1275 μm in depth. The probe was first coated in DiI stain in order to determine penetration location post-recording. The probe was then inserted at a 30° angle with respect to the vertical such that the probe was perpendicular to the surface of the barrel cortex in the left hemisphere. Recordings were acquired at 25 kHz and band-pass filtered above 300 Hz. Synapse software (TDT) was used to monitor activity on a TDT electrophysiological platform consisting of the PZ2 pre-amplifier and the RZ2 BioAmp Processor. All recordings were made while blind to genotype.

### Whisker stimulation

Nine whiskers were inserted into piezoelectric stimulators, generally by inserting the whisker that elicited the strongest response across all 64 channels in the middle of the array (Figure 1A) ^8,45^. A custom TDT program stimulated whiskers at different velocities. Each piezoelectric deflection was a ramp-hold-return (4 ms – 100 ms – 4 ms). To obtain a velocity response curve (VRC), whiskers were stimulated at 0, 65, 195, 326, 456, 587, 797 degrees per second, with varying amplitude and velocity throughout the recording period. 25 repetitions of each stimulation combination were recorded at 2-second interstimulus intervals, interleaving the deflections of whiskers and velocities. Spontaneous firing rate (no whisker stimulation) was measured in the 1 second before whisker deflection.

### Histology

Mice were euthanized with high dose isoflurane and cervical dislocation. The brain was isolated, cut at an approximately 30° angle, and sectioned on a vibratome with 225 μm sections. Sections were mounted and viewed under a Keyence microscope to determine probe penetration location (Figure 1B).

### Analysis

#### Spike sorting

A spike sorting algorithm was used to automatically sort waveforms, and units were then manually curated (both visually and with specific criteria) to identify single neurons. Recording files in TDT format were converted to a binary file using the TDTbin2mat function in Matlab (MathWorks). Data was then run through Kilosort 2.5 (spike sorting algorithm) on Ontologic’s platform (https://www.ontologic.ly/). The following changes were made to the preset parameters: threshold = [7 2], spike threshold = −5, sigma mask = 15, minimum firing rate = 0.1 ^46^. Manual curation of clusters then took place in Phy ^47^. During this process, clusters were either merged or separated based on waveform shape, cross- and auto-correlogram distributions, and template feature view. Noise waveforms (waveforms that did not have the characteristic shape of a neuron) were also discarded. All remaining clusters were then analyzed by downstream analysis (see below).

#### Single unit criteria

In order to ensure that clusters were truly single units, they had to pass certain criteria, based on past literature ^8,48,49^. We calculated the inter-spike interval (ISI) to determine the refractory period violation, which had to be < 1.5% in the 1 ms bin of the auto-correlogram. A mean firing rate was calculated throughout the length of the recording to ensure that units did not suddenly come into or out of the recording – less than 10% of the recording could be below 20% of the mean firing rate. In addition, < 20% of the missed spikes based on a Gaussian fit of the data was acceptable for a single unit. Finally, < 3% of the spike amplitude distribution could be below 11 μV. In order to differentiate putative excitatory and inhibitory neurons, we first graphed the trough to peak times for all units. We then fit a Gaussian Mixture Model to this distribution, which predicted an approximately 10% overlap between the two distributions (Figure 1C). For this study, we identified putative excitatory neurons as neurons with a half-width of more than 0.72 ms (Figure 1D). In addition, we re-ran all analyses after removing all units that have a half-width between 0.63 ms and .80 ms (the overlapping area), and found that the results remained similar, suggesting that the overlapping area contained mostly putative excitatory neurons.

#### Current source density (CSD)

CSD analysis was performed to elucidate the laminar position of electrodes on the silicon probe. First, the average local field potential (LFP) from stimulation-evoked responses of the best whisker (BW) was calculated. Then, the delta source method of inverse CSD (iCSD) was utilized to locate currents sinks and sources ^50^. The boundary between L4 and L5a was identified by the sharp change between current sink (L4) and current source (L5a). We assumed a width of 170 μm for L4 and 700 μm for L5/6 ^8,51,52^.

### Analyses conducted

The best whisker (BW) of the neuron was determined by performing a Wilcoxon rank sum test with p = 0.0056 (p = .05/9), to account for the 9 whiskers that were stimulated. The whisker that elicited the most significant response was deemed the BW for the neuron. The columnar whisker (CW) was defined as the whisker that corresponded to the barrel in which the probe was inserted. Probe penetrations that resulted in septal recordings were not included in the CW analysis, but were included for BW analysis.

Velocity response curves (VRCs) were generated using both the BW and CW stimulation for the neurons. The spontaneous firing rate for each neuron was subtracted from the evoked firing rate at each velocity, and a graph was produced to demonstrate how the firing rates changed over increasing velocities.

The proportion of neurons that significantly respond to a CW stimulation at the highest velocity was also quantified using a one-sided Wilcoxon rank sum test. Separate VRCs were generated for these neurons to determine if the CW-responsive neurons alter their response to whisker stimulation.

The minimum stimulation velocity needed to elicit a significant whisker-evoked response from a neuron was determined by using a one-sided Wilcoxon rank sum test at each stimulation velocity. This analysis was performed for all neurons that significantly responded to CW stimulation.

### Statistical analysis

A Two-Way ANOVA (mixed model) was used to determine if VRCs were significantly different when comparing two experimental conditions. Tukey’s HSD was then used to determine if there was a statistically significant difference at each velocity. In order to determine if there was a significant difference in the proportion of neurons that respond to CW stimulation, a z-score test was used. Finally, the Mann Whitney U test was utilized to compare the spontaneous firing rates.

## Code and Data Availability

All Matlab code (original and previously published) is available on Github (https://github.com/lakhanialishah/WhiskerAnalysis). Data for this paper will be available on The DANDI Archive (https://dandiarchive.org/dandiset/001171).

