## Supplemental Figures for "Whisker deprivation triggers a distinct form of cortical homeostatic plasticity that is impaired in the *Fmr1* KO"

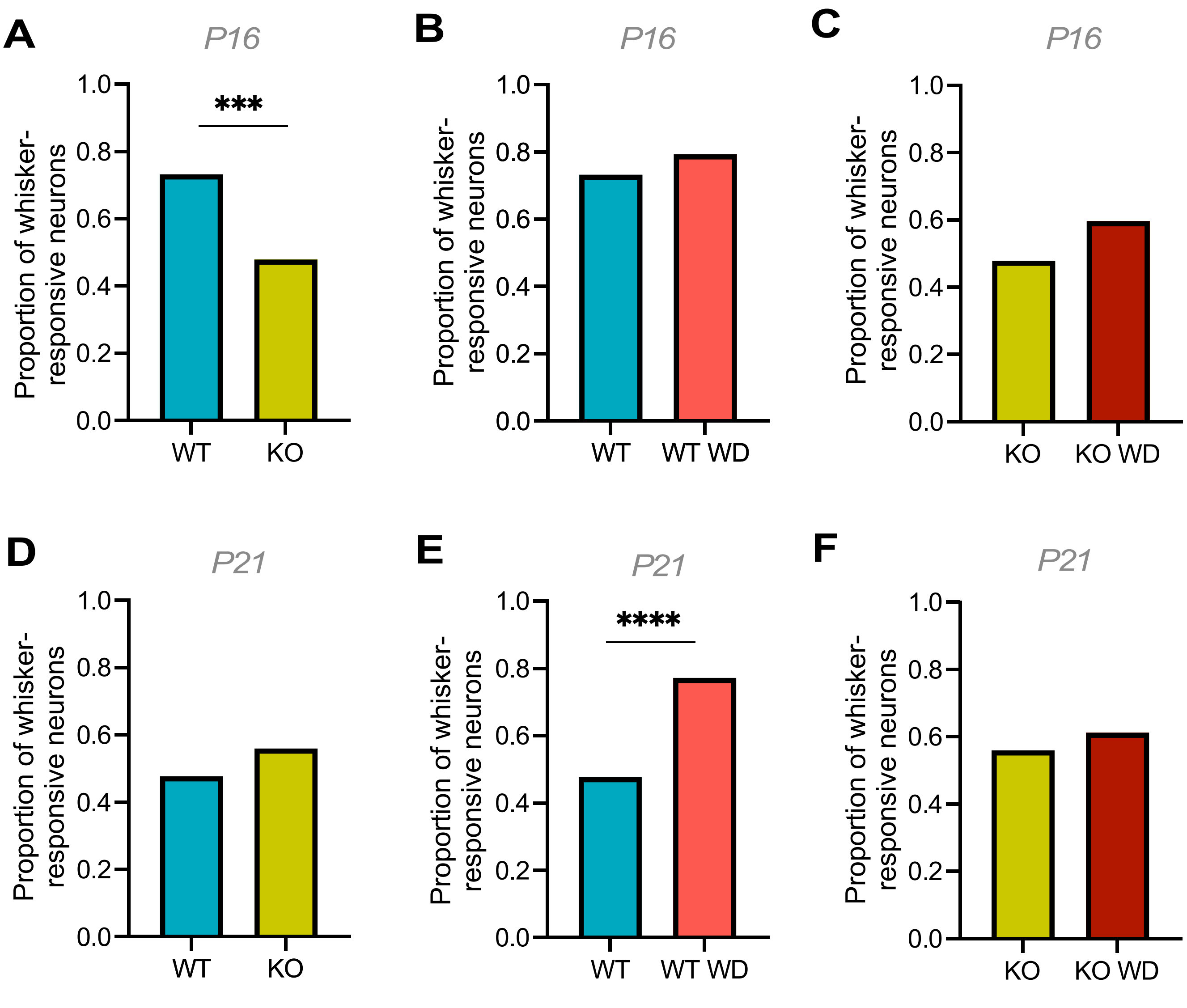


**Supplemental Figure 1. The proportion of whisker-responsive neurons in all experimental conditions, at baseline and following whisker deprivation.** A) P16 WT vs KO. B) P16 WT vs WT WD. C) P16 KO vs KO WD. D) P21 WT vs KO. E) P21 WT vs WT WD. F) P21 KO vs KO WD. *** $p < 0.001$, **** $p < 0.0001$.

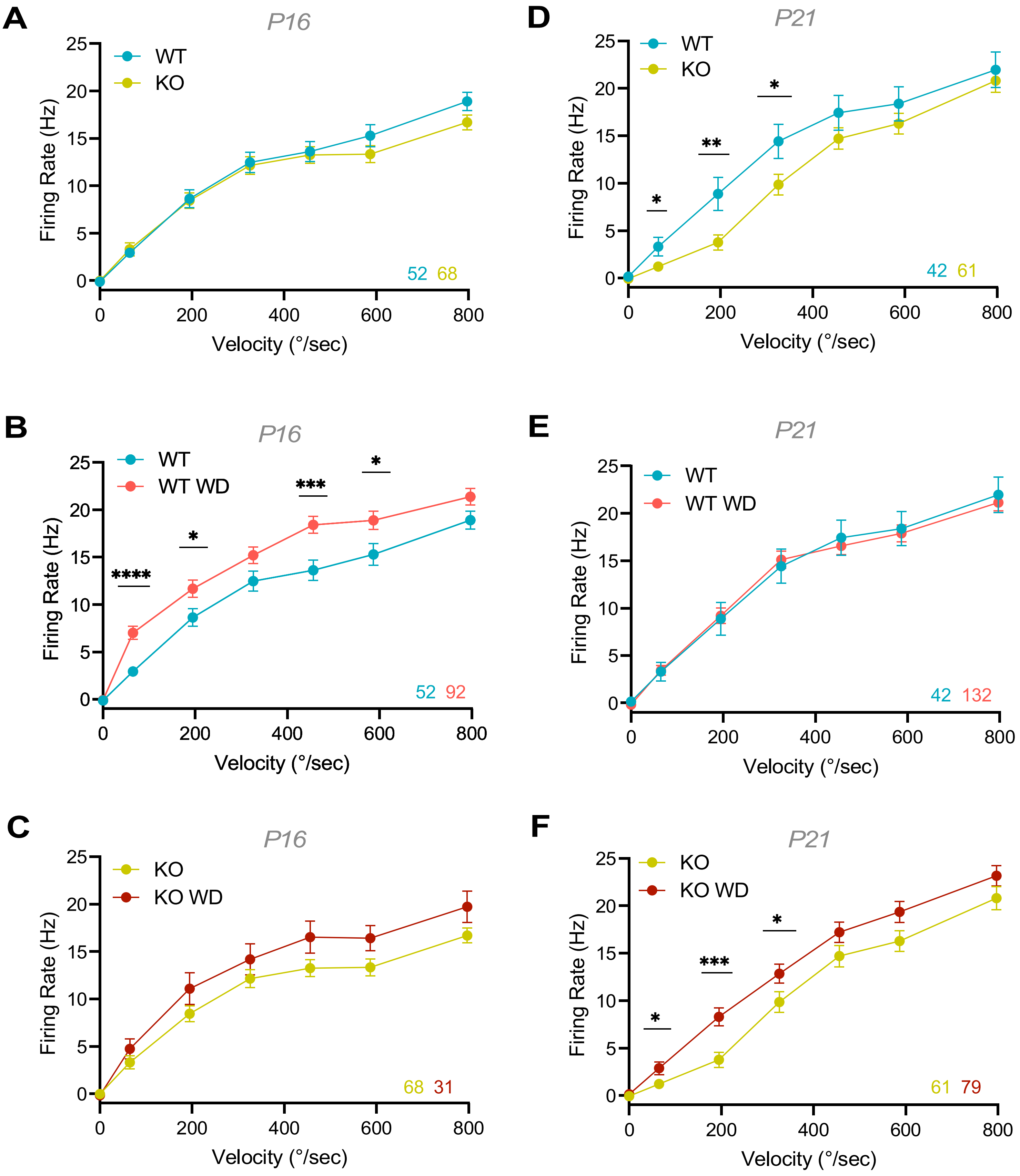


**Supplemental Figure 2. Best whisker (BW) velocity response curves (VRCs) of all experimental conditions.** A) P16 WT vs KO. B) P16 WT vs WT WD. C) P16 KO vs KO WD. D) P21 WT vs KO. E) P21 WT vs WT WD. F) P21 KO vs KO WD. Numbers at the bottom right are neurons per condition. Number of units for each condition is color-coded and shown at the bottom-right. * $p < 0.05$, ** $p < 0.01$, *** $p < 0.001$, **** $p < 0.0001$.
